# Sensory discrimination by isolated feeding tentacles in *Sanderia malayensis*

**DOI:** 10.1101/162867

**Authors:** Kazuo Mori, Matthew Sullivan, Joseph Ayers, Northeastern University, Marine Science Center, East Point, Nahant, MA 01908

## Abstract

Despite having only a diffuse neural network, tentacles of scyphomedusae exhibit a variety of distinct behavioral acts. One such behavior in tentacles is the capture and subsequent transfer of prey to the mouth. Interaction with prey consists of a variety of distinct stimuli—mechanical contact with the organism, chemical stimulation from the prey, and once captured by the tentacle, the weight of the food particles stretching the tentacle. By isolating and observing these individual stimuli in intact and isolated tentacles of the scyphomedusa, *Sanderia malayensis*, two separate reflexes have been distinguished. The tentacle feeding response observations suggest that the tentacle has two sensing networks, a bi-directional network for withdrawal and a proximally directed network for feeding.

**Summary Statement:** Two separate reflexes have been distinguished isolated tentacles of the scyphomedusa, *Sanderia malayensis*, a bi-directional network for withdrawal and a proximally directed network for feeding.

## Introduction

Jellyfish medusae perform a number of distinct behavioral acts, such as rhythmic swimming and food capture (Romanes 1880, Horridge 1953, Passano 1965). Two groups of jellyfish, hydromedusae and cubomedusa, those behaviors are controlled by their central nervous systems and peripheral neural net. Hydromedusae have a central nervous system that consists of two nerve rings to process sensory inputs and generate rhythms (Horridge 1955). The important neural elements for swim control in these animals are found in one of these nerve rings, the inner nerve ring. The *Aglantha digitale* produces an escape swimming response, in addition to slow spontaneous swimming contraction of the bell. Mackie and Meech (Mackie and Meech 1985) found that the motor giant neurons are able to produce two different types of spikes for these behaviors. They related different input events to discrete neural sub-systems. A comprehensive picture of the neuronal circuit was constructed by studying the synaptic inputs into the giant axons using extracellular recordings of the nerve rings in response to different stimuli (Mackie and Meech 1995, Mackie and Meech 1995, Mackie and Meech 2000, Mackie, Marx et al. 2003). Cubomedusae also have a central nervous system consists of the nerve ring and rhopalia those contain image forming eyes with lenses. A nerve ring connects (Garm, Ekstrom et al. 2006). Box jellyfishes, *Tripedalia cystophora* and *Chiropesella bronzie*, shows visually guided behavior suggesting that the CNS of these jellyfishes integrates visual information and innervate motor systems (Garm, O’Connor et al. 2007).

Recent studies on the Hydromedusae have used calcium imaging of genetically engineered Hydra to record the activity of all neurons (Dupre and Yuste 2017). They established that there are three distinct functional neuronal networks that mediate at least 4 different behavioral acts that have been established previously (Trembley 1744, Haug 1933, Ewer and Munro Fox 1947, Passano and McCullough 1963, Lasker, Syron et al. 1982).

A central nervous system or nerve rings on scyphomedusae have not been reported, but they exhibit swimming and feeding behavior similar to hydromedusae and cubomedusae that have a CNS. Pulsation rhythms are generated by any one of the rhopalia located on the bell margin (Romanes 1885). Scyphomedusae usually have 8 or 16 rhopalia and their pacemakers are responsible for initiating contractions. The rhythm of a rhopalium pacemaker can be reset by a current injection suggesting that an active pacemaker resets all others to avoid simultaneous initiations of bell contractions at different locations (Pantin and Dias 1952, Horridge 1959). There is no direct central axon tracts connecting rhopalia (Romanes 1885, Horridge 1954, Horridge 1956, Passano and McCullough 1963, Passano 2003), so signals are presumed to be conducted through the nerve net in the bell.

A rhopalium includes a statocyst, sensory epithelia, and ocelli (Schafer 1878, Martin 2002). The pacemaker is considered to be located in inner half of the rhopalium, however the exact location and circuitry of the pacemaker is unclear (Passano 1973, Nakanishi, Hartenstein et al. 2009). Rhopalia are connected to two kinds of nerve nets in the bell, the diffused nerve net (DNN) and the motor nerve net (MNN). Sensory inputs are sent to rhopalia through the DNN and an action potential from rhopalia that initiates bell contraction is transmitted to swimming muscles through the MNN (Romanes 1885, Horridge 1956). The DNN interacts with the MNN primarily at a rhopalium, and the DNN is also capable of enhancing MNN induced contractions (Romanes 1885, Horridge 1956, Passano and McCullough 1965, Passano 1973).

With the diffused nerve nets and rhopalia, scyphomedusae integrate and respond to environmental information. Arai reported that the scyphomedusa *Aurelia aurita* are attracted toward prey with which they are not in direct contact (Arai 1992). This result suggests that their swimming behavior is modified to produce a taxis, however, the precise network which underlies this modification has yet to be identified.

Capturing prey is an important event that can influence the pulsation rhythm of the bell. Some medusae, such as *Aurelia aurita,* have a number of short, fine tentacles on the bell margin to catch zooplankton. Other medusae, such as *Chrysaora hysoscella*, have fewer, longer tentacles and they feed on both zooplankton and larger prey such as fishes and other jellyfish. *Chrysaora* captures prey on a tentacle, and tangles it by the swimming movement of the medusa (Larson 1986). The tentacle then rapidly contracts, pulling the attached prey upward to the bell, where contact is made with an oral arm.

The tentacle of *Chrysaora* is composed of four basic layers: an outer epithelial layer, a layer of muscle fiber, a noncellular layer, called the mesoglia, and an endodermal layer which encloses the gastric canal (Burnett and Sutton 1969). Muscles are juxtaposed to the mesoglia (Perkins, Ramsey et al. 1971). A tentacle of *Chrysaora* contracts to a thirtieth of its relaxed length when prey is attached. Most of the length contraction arises from muscles crumpling into the mesoglia, only a small portion stems from actual shortening of the muscles themselves (Burnett and Sutton 1969, Perkins, Ramsey et al. 1971).

Anderson and co-workers (Anderson, Moosler et al. 1992) described immunohistochemically a nerve net in the outer epithelial layer of a *Chrysaora* tentacle. The region of nerve net stained on the subumbrella is considered a DNN (Anderson, Moosler et al. 1992), and it can be assumed that the stained nerve net would have a DNN like function. There might be another nerve net that is not immunoreactive to Antho-RFamide and innervates the motor system, like a MNN in the bell. Another nerve net has been stained in the endodermal layer; this would not innervate the motor system because mesoglia located between ectodermal and endodermal layers does not contain cells (Burnett and Sutton 1969). An extremely dense immunoreactive nerve net, the tentacular nerve ring, can be found in the ectoderm at their base (Anderson, Moosler et al. 1992).

*Sanderia malayensis*, is a tropical scyphomedusae that is classified into the same family to *Chrysaora* (Hargitt 1910). *Sanderia* rhythmically contracts its tentacle to keep the position of captured prey until an oral arm makes contact and contracts to transport this prey to the mouth. This behavior persists in isolated tentacles. This kind of rhythmic contraction has not been investigated and understanding the mechanism would be helpful to investigate signal processing by the neural net. Similarly to hydra (Dupre and Yuste 2017) it is possible to distinguish two different reflexes mediated by the neural net in isolated Sanderia tentacles. Through behavioral and electrophysiological experiments, we can distinguish individual sensory and motor components of these two tentacular responses of *Sanderia*.

## Materials and Methods

### Animals

Polyps of *Sanderia malayensis* were obtained from the New England Aquarium (Boston, MA, US). They were kept in circulating natural sea water at 22 °C. *Sanderia* polyps strobilated and released ephyrae at that temperature. *Sanderia* ephyrae were fed *Artemia* (Brine Shrimp Direct, UT, US) daily until their bell grew to 2 cm in diameter. Grown animals were fed chopped mussel or Aurelia every other day in addition to daily feedings of *Artemia*.

### Methods

All experiments were performed in ambient filtered sea water. A tentacle was isolated from medusa having bell diameters that ranged from 3 cm to 10 cm. Isolated tentacles were placed in a specifically designed perfusion chamber, 37 cm in height, 8 cm in width, and 3.5 cm in depth (Fig. 1). To record electrical activities, two Teflon coated silver wires were inserted into the gastric cavity separated axially by 4mm. We placed the tip of the distal (longer) electrode about 1 cm from the cut end of the relaxed tentacle. After the cut end was tied with a string, the tentacle was moved into the chamber, and was suspended by these electrode wires. When we did not use electrodes, the tentacle was suspended by a metal wire.

**Fig. 1.**
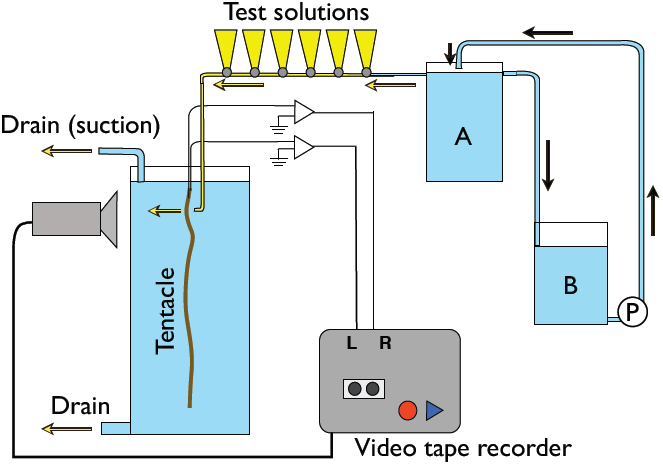
A diagram of the experimental setup. An isolated tentacle was suspended in a perfusion chamber with two metal electrodes inserted into the gastric canal. A gravity driven flow of sea water was continuously fed into the perfusion chamber (yellow arrow) from a holding tank (A). Sea water in the chamber was drained from the bottom, and the height was adjusted with a suction aspirator from water surface. The maximum level of the holding tank (A) is set by a drain near the top, that removes water in above this level to a holding tank (B). To ensure that the level in (A) is constant, sea water is continuously pumped from (B) to (A) using a pump (P). Tubes for test solutions were connected to the perfusion tubing with three way valves. Tentacle behaviors and electrical activities were recorded using a CCD camera on a videocassette recorder.

The recorded signals were amplified using an A-M systems differential AC amplifier. Tentacle behaviors were recorded using a CCD camera (SONY, XC-77) and recorded simultaneously with physiological signals on the audio tracks of a (SONY, GV-HD700) to provide synchrony between the behavioral and electrophysiological components. Signals were digitized as Quicktime Movies or digitized using an analog to digital converter (National Instruments, NI 6221) and acquired and processed with LabView software (National Instruments).

For chemical stimuli, we made a mussel juice by homogenizing and filtering mussels. We diluted the juice with sea water in a decade series. Amino acid solutions of glutamic acid, glutathione, asparagine, glycine, proline, valine, cysteine, methylglutathione and glutamine were made using sea water in concentrations of 10^−2^ M. Lucifer yellow (0.02 %) was dissolved in each test solution to visualize the stimulus making contact with the tentacle in the video signal. For glutamic acid and glutathione that were found to induce a response, 10^−5^, 10^−4^, 10^−3^, 10^−2^, 10^−1^ M solutions were prepared. During chemical stimulus experiments sea water was continuously fed into the perfusion chamber by gravity (Fig. 1). Sea water was drained from the bottom of the chamber and a vacuum aspiration was used to adjust the drain speed. Stimulus solutions were introduced into the chamber by temporarily switching the continuous flow of sea water to a flow of the stimulus solution.

Mechanical stimulus was given using a glass rod with a heat polished tip. We applied mechanical stimuli in three ways; 1) a slow touch that did not move the tentacle, 2) a quick touch, that resulted in tentacle motion, and 3) a rapid succession of quick touches.

In some experiments, tentacles were stretched by applying weight to the tentacle with square filter paper in sizes 2.5 mm, 5 mm, and 10 mm. The paper was introduced to the tentacle in the chamber with a tweezers and touched until the filter paper adhered to the tentacle, presumably through the release and attachment of nematocysts. After confirming attachment, the paper was released and allowed to stretch the tentacle.

Tentacle position was analyzed by tracking the position of a fiduciary mark on the tentacle, either a food particle, a cluster of nematocytes, or the filter paper stimulus, using digitizing tools (Hedrick 2008) in Matlab (The Math Works).

## Results

### 1. Feeding response in *Sanderia malayensis*

In the absence of stimuli, *Sanderia* medusa tentacles are in a relaxed state, freely floating as the organism swims. If contact is made with a large piece of food, such as mussel flesh, nematocysts are fired, capturing the prey. The tentacle then undergoes an immediate and large contraction, bringing the captured food toward the bell. Smaller prey, such as *`Artemia,`* will be captured in a similar manner, but does not induce an immediate tentacular contraction, although several such small food items will sum to eventually induce contraction in the tentacle. Concurrent with contraction, an oral arm swings toward the tentacle to transfer this food to the mouth. The food is then transferred from the tentacle to the oral arm. This process can be slow and during transfer the tentacle is observed to relax slightly followed by a subsequent contraction to bring the food back near the oral arm. Only once the food has been transferred does the tentacle relax completely.

### 2. Feeding response of isolated tentacles

A proximal repetitive contraction is observed in an isolated tentacle in response to a food stimulus. Tentacles were isolated from animals, either with or without the tentacular nerve ring. The tentacle was mounted in a seawater chamber and was not observed to contract in the absence of external stimuli. When a piece of flesh, either from a mussel or from a moon jelly, was attached to the tentacle, a proximal contraction was initiated immediately from the point of contact (Fig. 2). After fully contracting, subsequent contractions kept the food particle near the same, contracted position. This behavior is similar to that observed in intact animals, where the tentacle moves the food to an oral arm. After 45 seconds and several repeated contractions the tentacle relaxed fully and the food returned to its initial position. This demonstrates that the feeding reflex is endogenous to the peripheral neural net. The durations of this feeding response, from the first contraction to the full relax condition, were varied between preparations ranging from a minute to more than 15 min.

**Fig. 2.**
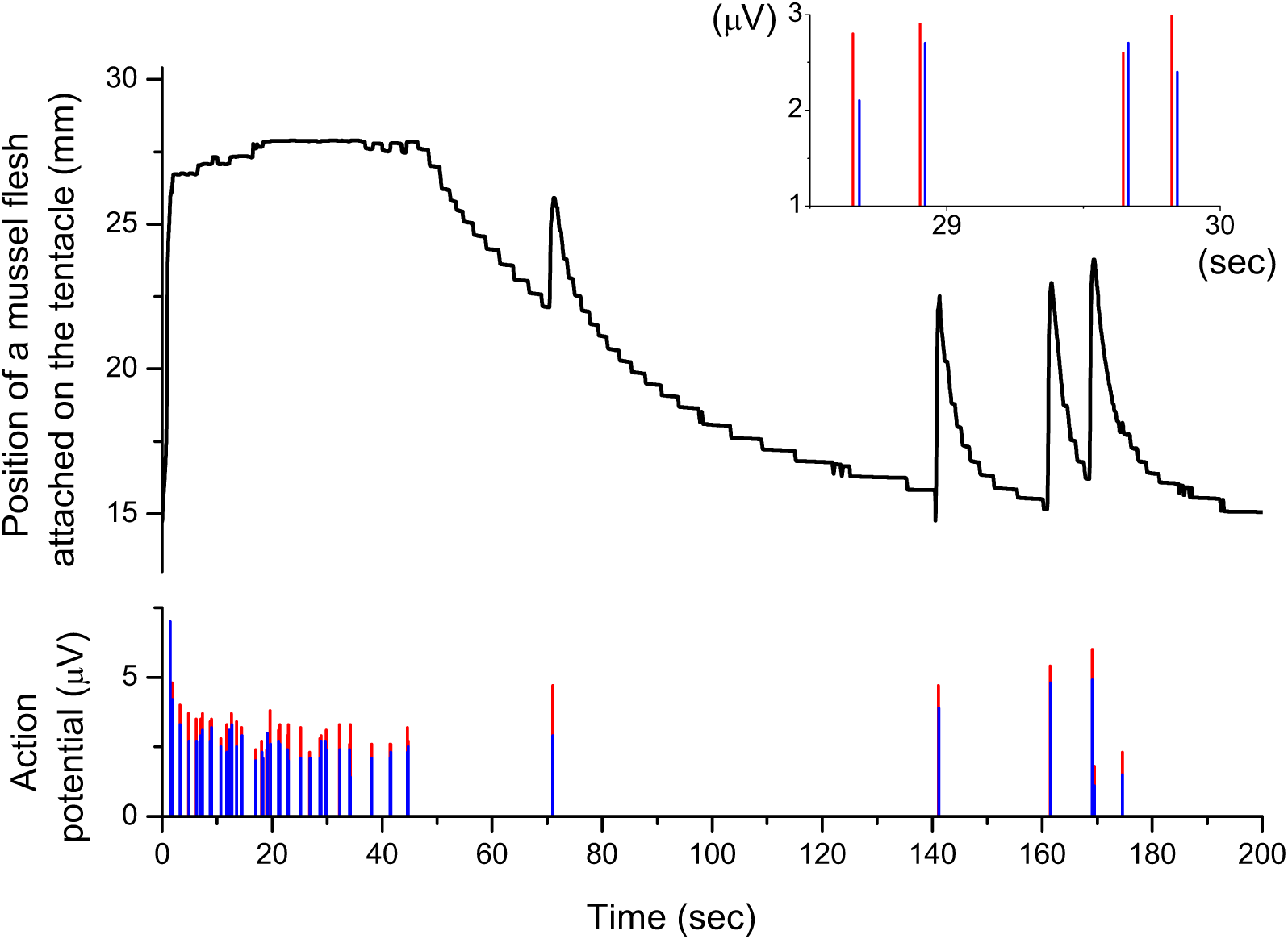
Tentacle extension and electrophysiological recording of an intact tentacle in response to a piece of mussel flesh. The tentacle extension is obtained by tracking an attached piece of mussel. Blue and red bars are two action potentials recorded with distal and proximal electrodes inserted in the gastric cavity of the tentacle. When a piece of mussel flesh was attached, at t = 0, it contracted proximally and it repeated small contractions to maintain the food position. Both action potentials followed contractions. The inset shows enlarged action potentials between 28 to 30 seconds. These contractions were initiated at the stimulated point on the tentacle and conducted proximally.

Concurrent with behavioral observations, electrophysiological measurements were made during the feeding response. Two wires inserted into the gastric cavity were attached at the base of the tentacle and spaced approximately 4 mm. Action potentials were observed coincident with every contraction (Fig. 2). Response from the distal electrode always preceded response from the proximal electrode, indicating the signal was travelling up from the tentacle (Fig. 2 inset).

A tentacular nerve ring that is located at the base of tentacle and is an extremely dense nerve net has been reported in two species *Chrysaora* and *Cyanea* those are classified into different families (Anderson, Moosler et al. 1992). It is assumed that *Sanderia*, classified into the same family as *Chrysaora*, also has a tentacular nerve ring. The feeding responses were observed on both of the tentacles prepared with and without the base that is expected to contain a tentacular nerve ring. However, the response of tentacles attached to the tentacular nerve net was longer in duration than those of isolated tentacles. Figure 3 shows a comparison of response durations of the two preparations to moon jelly and mussel stimuli. The duration was obtained by measuring the time between the first contraction after the stimulus was applied and the last contraction before the tentacle relaxed. Because the results were varied on preparations between 10 sec and 200 sec, the duration of the response an isolated tentacle was normalized with the duration of that of an intact tentacle isolated from the same animal. Then, the average was obtained with four experiments for each stimulus. Response durations of isolated tentacles were 25 % shorter than intact tentacles for moon jelly stimulus, and were 50 % shorter for mussel stimulus. This result suggests that the tentacular nerve ring may modulate the feeding response but is not necessary.

**Fig. 3.**
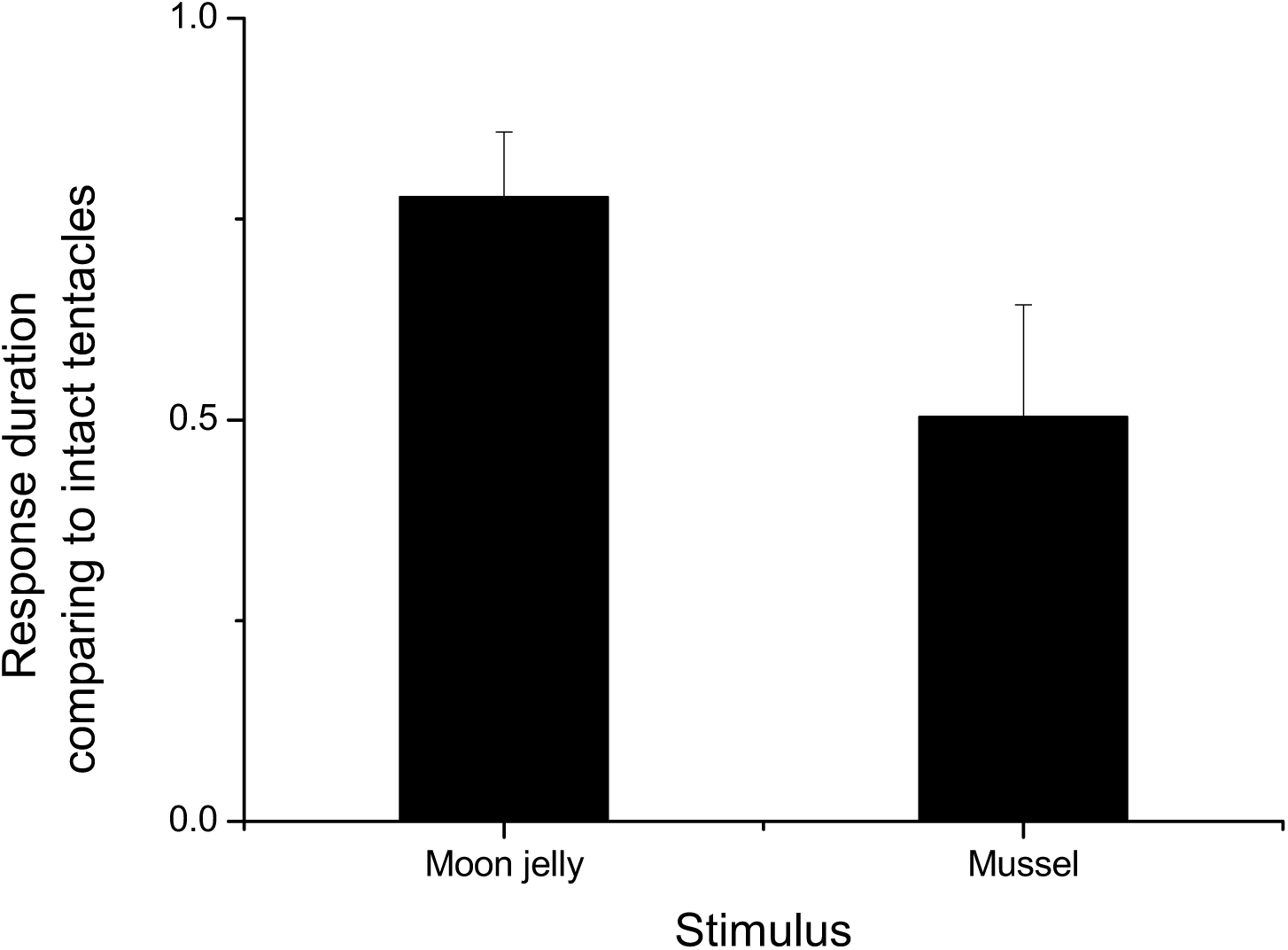
Comparison of response durations of isolated tentacles and intact tentacles. The response durations of isolated tentacles to moon jelly and mussel flesh are normalized by those of intact tentacles containing a tentacular nerve ring obtained from the same animal. These results are obtained with four tentacles for each experiment. Response durations of isolated tentacles were 25 % shorter than intact tentacles to a moon jelly stimulus and 50 % shorter to a mussel stimulus. This indicates that tentacular nerve ring located at the base of tentacles modifies response, but is not necessary to induce a response.

### 3. Feeding response to chemical stimuli

A food stimulus is complicated and can elicit many possible sensory reactions. Understanding the underlying sensory response requires simplifying the complicated food stimulus into component parts and understanding the isolated behavior of these responses. The chemical response is likely to be important for signaling the presence of food. The responses to two different chemical stimuli are studied, filtered juice from a homogenized mussel and solutions of amino acids.

To isolate chemical effects from mechanical ones, the chemicals are introduced to the tentacle through a constant stream of water flowing past the tentacle. A steady stream of fresh seawater is directed toward the tentacle under observation. There is no response observed to the stream of water itself. An experiment is performed by switching a valve upstream of the seawater outlet to inject a controlled amount of the chemical under test (200 µl).

Tentacle response to undiluted mussel juice was the same as the response to food stimuli. The tentacle immediately started repetitive proximal contraction and relaxed when the perfusion solution was switched back to sea water. The probability of invoking a tentacle response was decreased by decreasing the concentration. Below a dilution of 10^−3^ no response is observed (Fig. 4). At a dilution of 10^−1^ 11 of 12 tentacles proximally contracted, but localized contraction was observed before the proximal contraction. At a dilution of 10^−2^ 4 of 12 tentacles proximally contracted after a localized contraction. This result suggests that chemical stimuli alone can induce a feeding response and it has a threshold between 10^−2^ and 10^−3^.

**Fig. 4.**
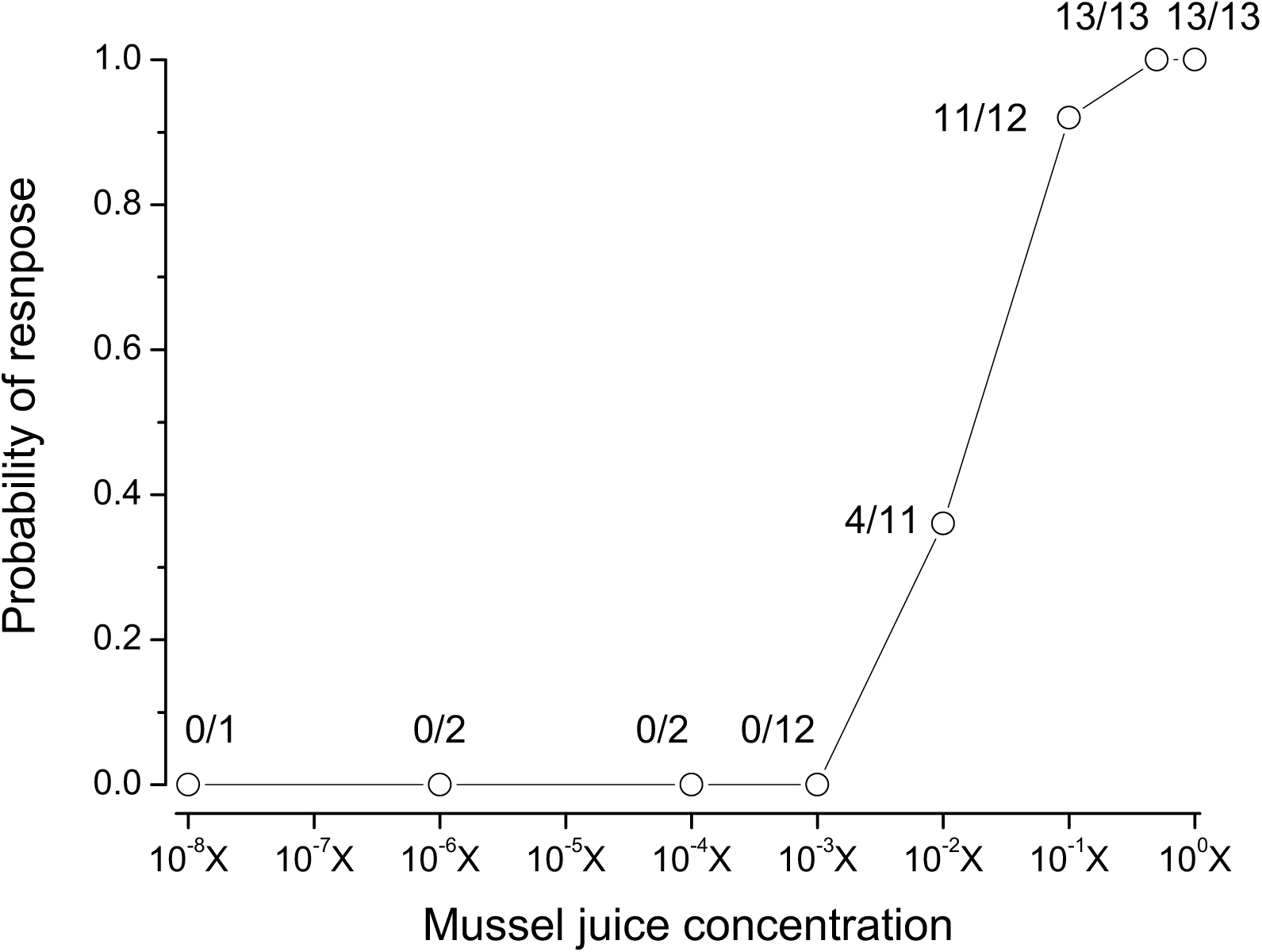
Dose response curve of tentacle to mussel juice. The number over the symbol indicates actual number of tentacles that responded to mussel juice and the number of tentacle tested. No tentacles responded at or below 10^−3^ times diluted mussel juice.

Comparison between the initiation of the first contraction at the stimulated point to 100 % mussel juice and the first measured electrical response recorded at the base of the tentacle allows for measurement of the signal delay. By adjusting the distance between the stimulation and observation points, the conduction velocity can be measured. The delay was increased linearly with distance, yielding a conduction velocity of 11 cm/sec (Fig. 5). We also used nine amino acids, glutamic acid, glutathione, glutamine, proline, asparagine, valine, methylglutathione, aspartic acid, and cysteine with concentrations of 10^−2^ M. In these amino acids, glutamic acid and glutathione could induce localized contraction. To investigate the detail of the responses to amino acids, we used 10^−1^ to 10^−5^ M glutamic acid and glutathione. No tentacles responded to these amino acids below 10^−3^ M. Tentacles responded to these amino acids at concentrations higher than 10^−2^ M with localized contraction. The response to glutathione was also localized contraction, but 3 of 6 tentacles proximally contracted to 10^−1^ M solution similar to the response to mussel juice. This result demonstrates that a single amino acid can excite the chemical sensory system to induce the feeding response of tentacles.

**Fig. 5.**
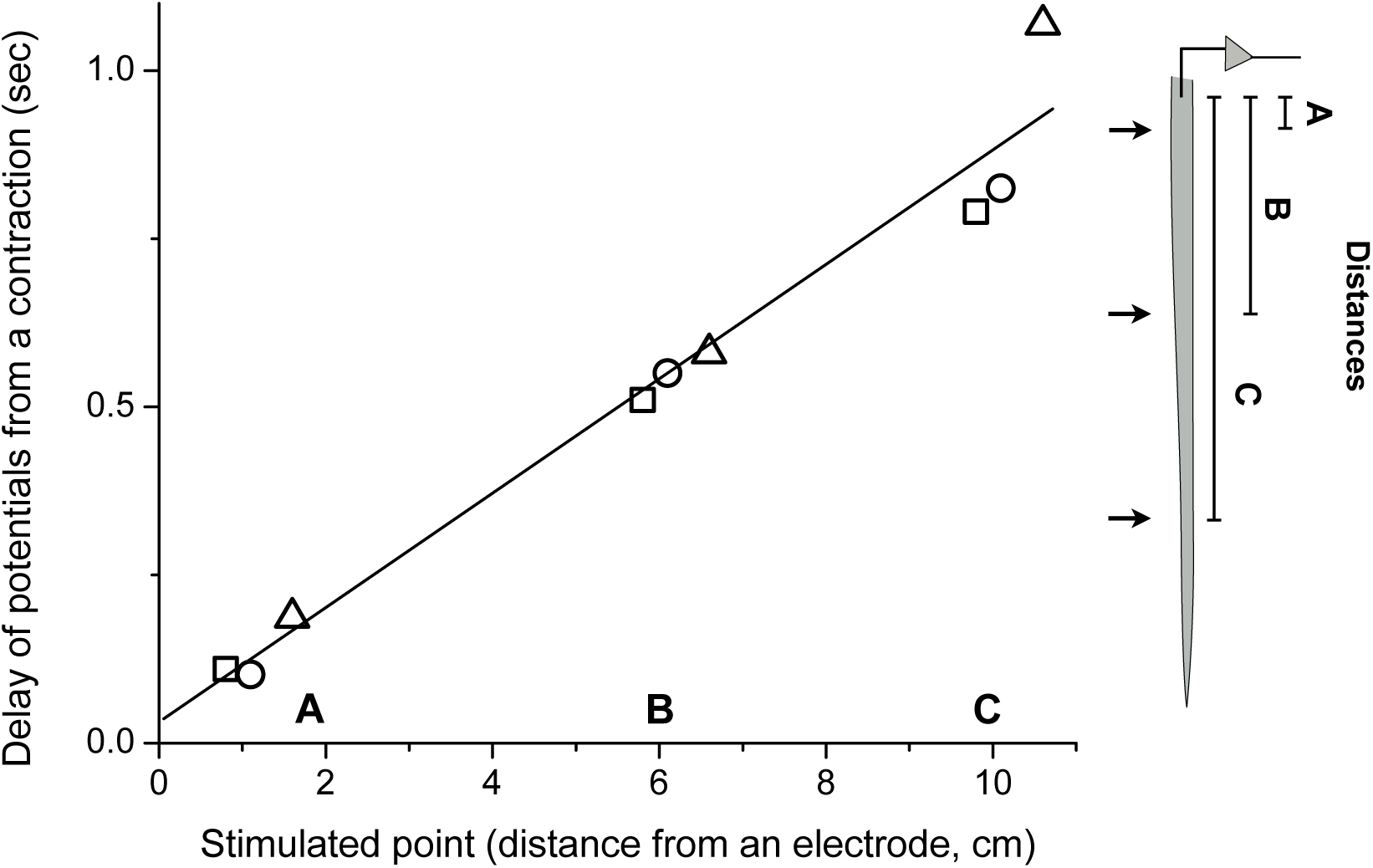
Delays of the first action potentials following the first large contraction to mussel juice applied to three points on a tentacle (right). This was obtained from three tentacles, and each symbol shows individual tentacle. The delay increased linearly by increasing the distance, and the conduction velocity was 11 cm/sec.

### 4. Mechanical stimuli induce a different response

Feeding is initiated by touching prey and mechanical stimuli are expected to be important for generating feeding behavior. We used a 1mm diameter heat polished glass rod as a mechanical stimulus by careful touching the tip of the glass rod to an isolated tentacle.

Four types of response, which we termed localized, distal, proximal and full contractions, could be induced by touching (Fig. 6). All tentacles tested locally contracted in response to slow contact (such that the tentacle remained stationary) (Fig. 6A). Tentacles sometimes adhered to the glass rod suggesting that nematocytes were discharged in response to this stimulus. With stronger touching (such that the tentacle was swung away from the point of contact) or repeated touching, distal, proximal or whole contractions could be induced (Fig 6 B, C, and D). However, in no cases were repetitive proximal contractions observed, as seen during the feeding response. Compared to chemical stimuli, the mechanical stimulus duration is short, less than 1 sec, and this short duration might be the reason for the absence of a repeated contraction. Experiments were also performed with constant mechanical stimulation using a glass rod. This stimulation induced a localized contraction just after a glass rod touch, but there was immediate relaxation and no repetitive contractions were observed. These results indicate that mechanoreception is not solely responsible for generating the feeding response.

**Fig. 6.**
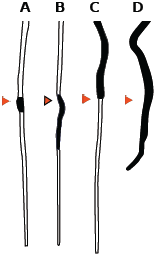
Tentacle contraction patterns: localized (A), distal (B), proximal (C) and whole contractions (D) induced by slow (A), fast (B and C) and fast and repeating touching (D) with a glass rod, respectively. Red rectangles indicate the stimulated points on tentacles and contracted areas on tentacles shown as black areas.

### 5. Stretch stimulus induced repetitive proximal contraction

Repetitive tentacle contraction was not induced by a single small prey, for example *Artemia*, but could be by the capture of larger prey or of multiple *Artemia* on the same tentacle. Therefore, there appears to be a response to tentacle stretching that is independent of the touch response. To address this sensory mechanism, we used a piece of filter paper that is negatively buoyant and stretches the tentacle when attached, but does not contain any proteins or amino acids.

To control for the purely tactile response, we first investigated the influence of the paper by touching the tentacle at a fixed position, with the filter paper supported by a thread. The response to this stimulation was a localized contraction and immediate relaxation (Fig. 7A), similar to the response to constant mechanical stimulus with a glass rod.

**Fig. 7.**
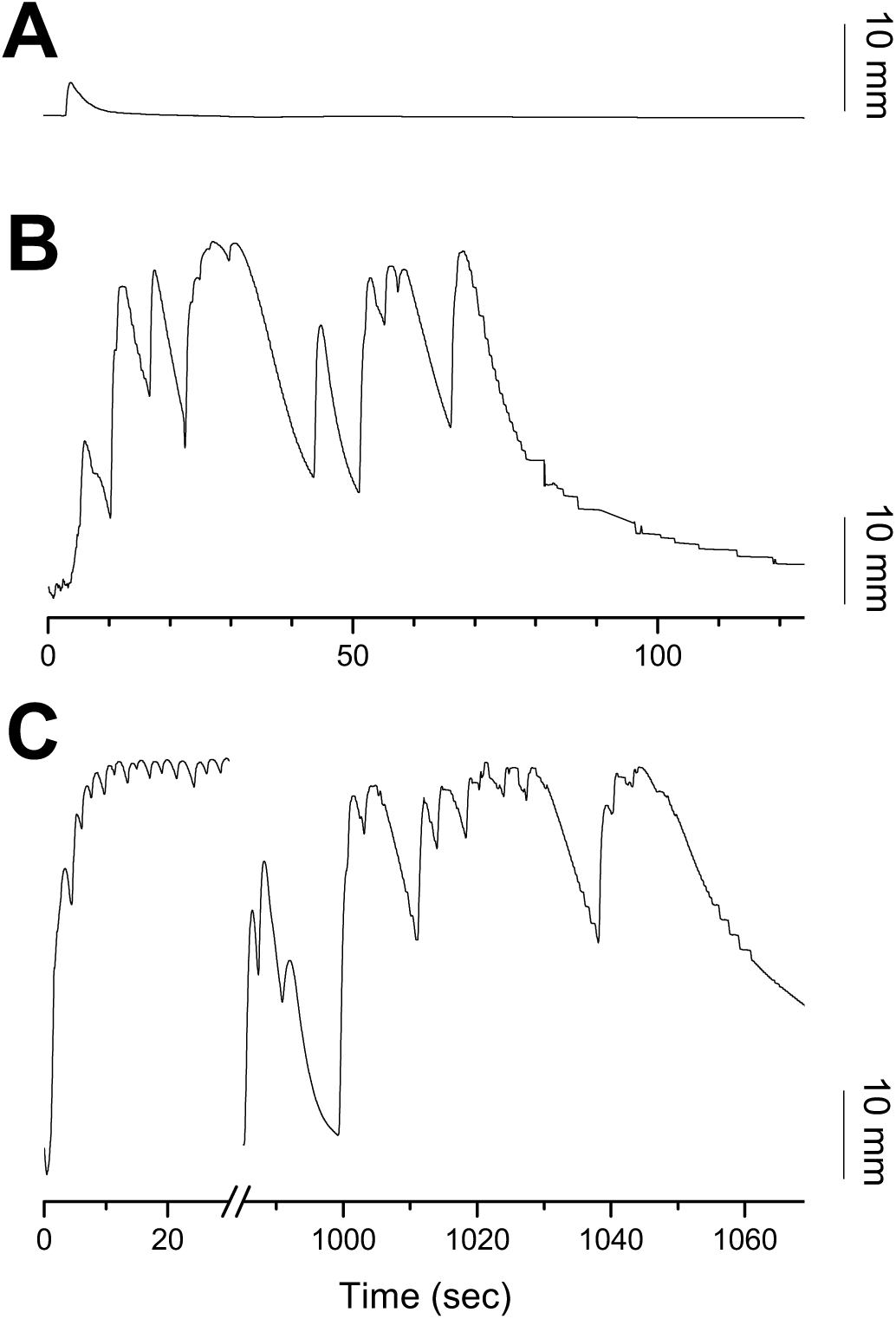
Contraction patterns in response to paper and food stimuli on a tentacle. (A) Response to touch stimulus with a vertically suspended piece of paper. There is local contraction followed by immediate relaxation. (B) Response to a piece of paper attached directly to the tentacle. Repetitive proximal contraction was recorded. (C) Response to a piece of mussel shown in two elapsed time regions, 0 to 20 sec and 990 to 1070 sec at the same scale as in A and B. The tentacle repeated contraction with faster frequency and smaller amplitudes just after it was stimulated than the late repetitive contraction. This late response shows the same qualitative features found in attachment of paper.

With mechanical stimulation, the filter paper would often become attached to the tentacle, likely through a nematocyte. To induce a stretch stimulus, the paper was released after attachment, causing it to sink due to gravity. After some strain of the tentacle, a repetitive proximal contraction was observed, originating from the contacted point (Fig. 7B). This response could be induced by attaching filter paper to any point on tentacle. Compared to the food response, the frequency of the contraction of the paper response was lower and the amplitude was bigger. Figure 7C shows the beginning and end of the response to a piece of mussel, with remarkable similarity between the late stage response to a food particle and the response to filter paper. This suggests that the food particle first stimulates the chemical and stretch responses, and only later does a stretch response induce repetitive contractions.

Like a chemical stimulus, the stretch stimulus induces a repetitive proximal contraction. The two stimuli have different regions of excitation, chemical stimuli being localized on the applied point while stretching occurs from the point of contact proximally to the fixed base of the tentacle. To address the directionality of this response, we performed stretch experiments with a gravitationally inverted tentacle. A paper stimulus placed on this inverted tentacle stretched it distally from the point of contact, rather than the usual proximal stretching. Even with this distal stimulus, repetitive proximal contractions were induced, suggesting that this directionality stems from the organism and not from the orientation relative to gravity.

## Discussion

An intact scyphomedusa, *Sanderia malayensis*, keeps swinging tentacles by bell pulsations during it swims. Prey is captured on a tentacle and is brought to near the bell by a tentacle proximal contraction, and the tentacle then repeats contraction to keep the prey position near the bell until it transfers the prey to an oral arm. We found that isolated tentacles could produce this feeding response suggesting that this behavior would be generated with simple and primitive nerve nets in the tentacle. This study of *Sanderia* tentacle demonstrates that the isolated tentacle nerve net can discriminate between chemical and tactile stimuli and generate at least two different behavioral acts.

Contractions on an isolated tentacle could be initiated by mechanical, chemical and stretch stimuli, but these different stimuli led to different types of response. Since prey must be touched on a tentacle to induce feeding response, touching is considered to be one of the triggers of the response. However, mechanical stimuli did not induce the feeding response on tentacles. To a mechanical stimulus tentacle contracted locally, distally, proximally or whole length (Fig. 6). After these contractions, the tentacle relaxed immediately and did not repeat contraction. Horridge (Horridge 1955) reported a tentacle response to mechanical stimuli on a hydromedusa, *Geryonia proboscidalis*. Tentacles on an intact animal proximally contracted to a mechanical stimulus while an isolated tentacle contracted locally to gentle stimulus or the whole tentacle contracted to a strong stimulus. This indicates that the CNS of *Geryonia*, that consists of inner and outer nerve rings in the bell, strongly innervate tentacular nervous system. However, in scyphomedusa that does not have such CNS (Satterlie 2011), intact *Sanderia* exhibited variety of temporal responses by mechanical stimuli those are the same to the responses of isolated tentacles. This result suggests that tentacular nervous system of *Sanderia* is not strongly innervated by nerve nets in the bell. This variety of temporal responses of tentacles by mechanical stimulation might function to allow escape from predators or to reject non-food objects, but not trigger the feeding response.

In cnidarians, chemical stimuli are well known as feeding response activators and a specific amino acid works to induce a specific step of feeding behavior in some species. In sea anemone, for example, proline triggers a food capture response while glutathione triggers ingestion (Lindstedt 1971). In scyphomedusa, polyps of *Chrysaora* exhibit feeding behaviors in response to various amino acids and peptides, and a specific feeding step for each amino acid such as contraction or writhing was also observed in excised tentacles (Loeb and Blanquet 1973). For a *Sanderia* medusa tentacle, filtered mussel juice and nine amino acids were used to investigate the roles of chemosensory system on the feeding response. The mussel juice induced repetitive proximal contraction, but very high concentration, more than 1 % is required. Glutamic acid and gluthatione induced a localized contraction to 10^−2^ M amino acids. To 10^−1^ M of glutamic acid only localized contraction was produced, while 10^−1^ M of glutathione produced proximal contractions in addition to a localized contraction on 3 of 6 tentacles. This low sensitivity to chemical stimulus might be useful to avoid responding to a bleeding prey located too far from the tentacle, but responses of the chemosensory system on a tentacle can trigger the feeding response with strong stimuli.

Because a few *Artemia* captured on a tentacle did not evoke a tentacle contraction but larger prey was able to, it is considered that tension on a tentacle would be required. Our paper stimulus induced repetitive proximal contraction from the stimulated point when allowed to stretch the tentacle, while touching with a fixed paper only caused a temporal localized contraction. These results indicate that a stretch receptor is a trigger and a modulator of the feeding response allowing it to persist until the prey is transferred to the oral arm. Unlike chemical and mechanical stimuli, which are applied on a stimulated point of the tentacle, tension in the tentacle is usually present proximally from the point of attachment. Despite this broad area of stimulation, the response of the tentacle seems to be initiated at the point of contraction and proceed proximally, even if the stimulus has been applied distally by inverting the tentacle. How stretch receptors located on the proximal or distal stimulated area is unclear, and to investigate the mechanism more detailed experiments are required.

Rhythmic contraction could be evoked on any part of tentacle through the application of a food, stimulus, or stretch stimuli. This rhythmic contraction arises from interplay between prey weight continuously pulling the tentacle down while muscle response intermittently moves the food back up. Removing the stimulus results in cessation of rhythmic contraction, indicating the necessity of continuous chemical or stretch stimuli to induce repeated contraction. As the tentacle habituated to the stimulus, the frequency of contraction decreased, but this decreased frequency was accompanied by increased amplitude to ensure the food was raised to its previous location (Fig 2 and 7C). To generate rhythmic patterns, medusa has rhopalia on the bell edge contains a pacemaker that initiates a bell contraction and the rate of firing of individual rhopalial pacemaker is variable and can be directly influenced by sensory inputs, which presumably modify a baseline discharge rate of the pacemaker (Romanes 1885, Horridge 1956, Passano and McCullough 1965, Passano 1973). However, such rhoparium like organs have not been found on the tentacle, and our results indicate that the rhythmic pattern is generated in the tentacle. The response duration of the preparation contained tentacular nerve ring was significantly longer than preparations those did not contain the tentacular nerve ring (Fig. 3), but the response pattern was the same, repetitive intrinsic proximal contraction of which the tentacular nerve ring could be an extrinsic modulator. The cellular nature of the pacemakers of tentacle nerve nets are not known, but the sensory signals would be integrated at interneurons which must be located along the tentacle, and the firing rate would be modified by sensory input levels from both chemical and stretch receptors.

Responses to chemical and stretch stimuli signals were always conducted proximally, suggesting these responses are connected to a proximally conducting network. This is different from mechanical stimuli that could induce a response in both directions suggesting that existence of two nervous systems in the tentacle. The bell contains two nerve nets, a motor nerve net (MNN) and a diffuse nerve net (DNN), both of which consist of bidirectional synapses between neurons (Horridge and Mackay 1962, Horridge, Chapman et al. 1962). In analogy to the bell nervous system, there might be two nerve nets on the tentacle: one for feeding response and polarized to conduct proximally (feeding nerve net) and one for escape or rejection behaviors which conducts either direction depending on stimulus conditions (withdrawal nerve net). The feeding nerve net would be comparable to MNN that innervate muscles directly and the withdrawal nerve net is similar to the DNN that modulates muscle activities (Romanes 1885, Horridge 1956, Passano and McCullough 1965, Passano 1973). However, the bell MNN is not polarized to conduct signals (Anderson 1985, Anderson and Spencer 1989), and this is a major difference between the proposed polarized feeding nerve net and the MNN. This suggests that the feeding nerve net would have polarized synaptic projections, however, more detailed histological and physiological information on the tentacle nervous system are required.

From the perspective of function, it makes sense that stimuli distinctly associated with food, those of chemical and stretch detection, produce different responses than touch stimuli associated with both food and predation. The rapid withdrawal behavior associated with full distal and proximal contraction is distinctly different than the repeating and more sustained feeding behavior of raising capture prey for transfer to the oral arms. On our preliminary experiments, an isolated oral arm could also be induced a feeding response by a food particle suggesting that the feeding behavior of the oral am is also generated by endogenous nervous activities. These results suggest that the feeding response of *Sanderia* medusa consists of two endogenous processing systems located in a tentacle and an oral arm system coordinated through nerve nets in the bell.

## Acknowledgements

We thank Martin Poitsch for support and valuable feedback.

## Competing interests

The authors declare no competing or financial interests.

## Author contributions

Conceptualization: K.M., M.S., J.A.; Methodology: K.M., J.A.; Formal analysis: K.M., M.S., J.A.; Investigation: K.M., M.S.; Resources: J.A.; Writing - original draft: K.M., M.S; Writing -review & editing: K.M., M.S., J.A.; Supervision: J.A.; Project administration: J.A.; Funding acquisition: J.A.

## Funding

Supported by Schlumberger-Doll Research, Cambridge, MA.

